# A role for N6-methyldeoxyadenosine in *C. elegans* mitochondrial genome regulation

**DOI:** 10.1101/2023.03.27.534452

**Authors:** Lantana K. Grub, James P. Held, Tyler J. Hansen, Samantha H. Schaffner, Marleigh R. Canter, Evi M. Malagise, Maulik R. Patel

**Affiliations:** Department of Biological Sciences, Vanderbilt University, Nashville, TN; Department of Biochemistry, Vanderbilt University, Nashville, TN; Department of Cell and Developmental Biology, Vanderbilt University School of Medicine, Nashville, TN; Diabetes Research and Training Center, Vanderbilt University School of Medicine, Nashville, TN

**Keywords:** Mitochondria, mtDNA, epigenetics, 6mA

## Abstract

Epigenetic modifications provide powerful means for transmitting information from parent to progeny. As a maternally inherited genome that encodes essential components of the electron transport chain, the mitochondrial genome (mtDNA) is ideally positioned to serve as a conduit for the transgenerational transmission of metabolic information. Here, we provide evidence that mtDNA of *C. elegans* contains the epigenetic mark N6-methyldeoxyadenosine (6mA). Bioinformatic analysis of SMRT sequencing data and methylated DNA IP sequencing data reveal that *C. elegans* mtDNA is methylated at high levels in a site-specific manner. We further confirmed that mtDNA contains 6mA by leveraging highly specific anti-6mA antibodies. Additionally, we find that mtDNA methylation is dynamically regulated in response to antimycin, a mitochondrial stressor. Further, 6mA is increased in *nmad-1* mutants and is accompanied by a significant decrease in mtDNA copy number. Our discovery paves the way for future studies to investigate the regulation and inheritance of mitochondrial epigenetics.

## Introduction

Epigenetic modifications are a powerful tool for genomic and transcriptomic regulation. One of the most common DNA modifications is N6-methyldeoxyadenosine (6mA). 6mA is the most abundant methylation mark in prokaryotes and is present in multiple clades of bacteria. This high level of conservation suggests it has an essential role in cell regulation ^1^. It has been well studied in the context of self-sensing restriction modifications ^2^ and also has critical roles in prokaryotic DNA replication and chromosome segregation ^3^, transcription ^4^, DNA repair ^5^, and cell cycle regulation in some species ^6^.

In addition to its role in prokaryotes, recently, it was discovered that 6mA is an epigenetic mark in lower-level eukaryotes. 6mA is highly abundant in the green algae *Chlamydomonas reinhardtii* nuclear genome and presents with a bi-modal clustering near transcription start sites and marks linker regions between nucleosomes ^7^. *Chlamydomonas* 6mA sites are significantly enriched at GATC sequence motifs, which is analogous to bacterial epigenetics. 6mA is also found in early-diverging fungi where it is concentrated in promoter regions of protein coding genes and is correlated with high gene expression ^8^.

6mA is also an important genome regulator in some complex, multicellular animals. *Drosophila* have high levels of nuclear 6mA across transposons in embryos which is depleted over the course of development ^9^. Nuclear 6mA and its regulatory enzymes have also been described in *Caenorhabditis elegans*. 6mA of the nuclear genome is not associated with specific gene features in *C. elegans* ^10^, but 6mA can be transgenerationally inherited and can serve as a powerful form of stress adaptation ^11^. 6mA mediates stress response at least in part by marking sites for the transcription factor ATFS-1 to bind to activate the mitochondrial unfolded protein response (UPR^mt^).

New roles continue to emerge for 6mA, including a potential role in regulation of mitochondria. Because of their ancestral link to alpha proteobacteria, mitochondria retain their own small, but essential, genome (mtDNA). Given the prokaryotic origin of mitochondria and the capacity for eukaryotic cells to facilitate 6mA methylation, are the genomes of mitochondria 6mA modified? Indeed, recent publications suggest that mtDNA from human cell culture contains 6mA ^12, 13^. However, the experimental strategies employed in these studies have limitations in assessing 6mA methylation ^14^. It is therefore necessary to establish mtDNA methylation using a variety of complementary approaches that supplement the limitation of each experiment.

We first aimed to convincingly determine if mtDNA is 6mA methylated. Secondly, if mtDNA is methylated, we reasoned that it would be most beneficial to characterize it in an *in vivo* system in which inter- and transgenerational experiments can be conducted. To answer both questions, we chose to determine if mtDNA is methylated in *C. elegans*. *C. elegans* is a powerful genetic model with a short generation time making it an ideal system for studying mitochondrial epigenetics.

We sought to generate *in vivo* evidence of 6mA methylation in the *C. elegans* mitochondrial genome. First, we analyzed two publicly available data sets that originally assessed 6mA levels in *C. elegans* nuclear DNA but did not analyze mtDNA. We find high 6mA levels in the mitochondrial genome in both a Single Molecule Real-Time (SMRT) sequencing data set ^10^ and significant mtDNA enrichment in a 6mA methylated DNA IP (MeDIP) data set ^11^. We then directly assessed 6mA of the mitochondrial genome using two complementary biochemical assays. Dot blot for 6mA in mitochondrial enriched DNA preparations and immunoprecipitation for mtDNA using an anti-6mA antibody, both of which show mtDNA 6mA methylation. Moreover, mtDNA could be competitively eluted from 6mA antibodies with 6mA containing synthetic oligonucleotides but not with non-methylated oligonucleotides further supporting the specificity of our antibody to detect 6mA on mtDNA. Using the SMRT sequencing data, we also characterized mtDNA 6mA patterns at single base resolution and show that 6mA methylation patterns are non-random, suggesting that 6mA may provide regulatory function. Finally, to interrogate the potential physiological relevance of mtDNA 6mA, we assessed mtDNA 6mA in response to mitochondrial stress. mtDNA 6mA increases in response to mitochondrial stress induced by treatment with the electron chain inhibiting compound antimycin suggesting 6mA can be dynamically regulated. Methylation levels also change in response to genetic perturbations. We find that knock out of NMAD-1, a demethylase domain containing protein, increases methylation and this correlates with a decrease in mtDNA copy number and mild but reproducible activation of the mitochondrial unfolded protein response (UPR^mt^). Combined, these data strongly support that *C. elegans* mtDNA is functionally 6mA methylated and provide one of the first characterization of an *in vivo* model for studying mtDNA epigenetics.

## Results

### SMRT sequencing and MeDIP data sets show high 6mA methylation in mitochondrial DNA relative to the nuclear genome

Recent advances in third generation sequencing technology have made high throughput study of mtDNA methylation possible. Techniques such as Single Molecule Real-Time (SMRT) Sequencing (PacBio) and Nanopore Sequencing (Oxford Nanopore) sequence native, long-read DNA in real time, providing both sequence information and epigenetic modification status by distinguishing between modified and unmodified bases ^15, 16^. Using a previously published SMRT sequencing data set, we found that mtDNA contains the epigenetic signature N6-methyldeoxyadenosine (6mA). 862 of the 10,514 adenines of the mitochondrial genome had a methylation prediction with p-value ≤ 0.001, which represents a 6mA:dA ratio of 8.2% (Figure 1A). This is approximately a 20-fold higher 6mA:dA ratio than that of nuclear chromosomes, which is 0.4% on average (Figure 1A). Next, given that there are more copies of mitochondrial DNA in cells than nuclear DNA, we asked if this increase in 6mA:dA ratio was due to higher coverage of the mitochondrial chromosome relative to nuclear chromosomes in the SMRT sequencing data. The unique 6mA sites of mtDNA had 320x coverage on average, whereas the nuclear chromosomes had an average of 63x coverage (Supplemental Figure 1A and B). This 5-fold increase in coverage does not independently account for the 20-fold higher 6mA:dA ratio.

**Figure 1.**
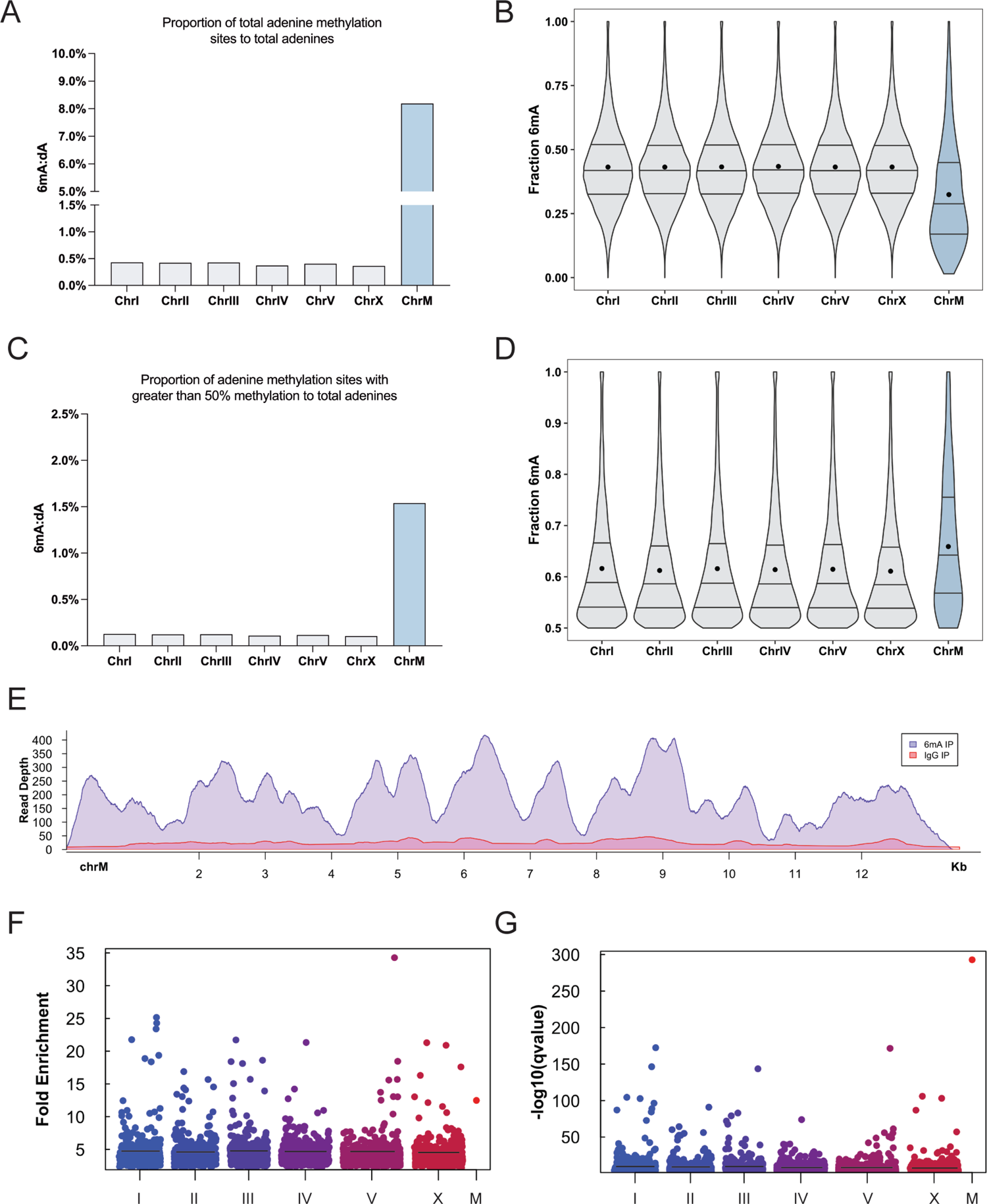
SMRT sequencing and MeDIP data sets show high 6mA methylation in mitochondrial DNA relative to the nuclear genome. (A) The ratio of total adenine methylation sites (6mA) with a p value ≤ 0.001 to total adenines (dA). Nuclear chromosomes shown in gray, and the mitochondrial chromosome in blue for Figure 1A-D. ChrI n=41,916, ChrII n=41,457, ChrIII n=37,943, ChrIV n=42,763, ChrV n=54,567, ChrX n=42,009, ChrM n=862. Means are shown. (B) The fraction of reads that are identified as methylated, from 0.0 to 1.0, for a unique adenine. Means (black dot) and distribution quartiles (lines) are shown. (C) The 6mA:dA ratio of sites with 50% methylation or greater. ChrI n=12,269, ChrII n=11,915, ChrIII n=11,028, ChrIV n=12,527, ChrV n=15,744, ChrX n=12,100, ChrM n=162. (D) The fraction of reads that are identified as methylated, from 0.5 to 1.0, for a unique adenine. Means (black dot) and distribution quartiles (lines) are shown. (E) MeDIP read depth throughout the mitochondrial chromosome. Control IgG IP shown in red, 6mA IP in purple. (F) The fold enrichment of the 6mA IP over the control IP for each significantly enriched sequence. The mean fold enrichment for each chromosome is shown (black line). (G) Unique sequences that are significantly enriched in the 6mA IP compared to the control using MACS2. The -log10(qvalue) for each enriched sequence is plotted as a single point. ChrI n=574, ChrII n=535, ChrIII n=513, ChrIV n=554, ChrV n=588, ChrX n=654, ChrM n=1. The mean -log10(qvalue) for each chromosome is shown (black line).

We then asked, what the incidence of methylation is at each unique site (i.e. for each methylated adenine site, what fraction of reads are methylated). The average methylation level at 6mA sites in the mitochondrial genome was 32% compared to an average methylation level of 42% in the nuclear chromosomes (Figure 1B). However, higher sequence coverage of mtDNA relative to nuclear DNA may result in greater ability to detect sites with lower levels of 6mA. This is supported by a significant correlation, Spearman’s rank correlation rho = −0.5668, between a higher sequence coverage and a smaller confidence interval of methylation percentage at a given site (Supplemental Figure 1C and D). To account for the difference in ability to detect 6mA at low levels, we filtered the data to include only sites with at least 50% or greater methylation fraction. After filtering, there was a 10-fold higher 6mA:dA ratio in mtDNA compared to the nuclear chromosomes (Figure 1C). Additionally, after filtering, the average methylation level at a 6mA site is similar in mtDNA and nuclear chromosomes, 65.9% and 61.3% respectively (Figure 1D).

To complement the SMRT sequencing data, we analyzed a methylated DNA immunoprecipitation (MeDIP) sequencing data set from Ma et al. that immunoprecipitated DNA with an anti-6mA antibody and then performed Illumina sequencing ^11^. This analysis further confirmed 6mA in mtDNA (Figure 1E). We compared the read depth of DNA in the control IgG vs 6mA IP. This identified sequences with a significantly increased read depth, indicating 6mA within that sequence. mtDNA was significantly enriched for 6mA compared to the non-specific IgG control. The methylation fold-enrichment for mtDNA was substantially higher than the average fold-enrichment in the nuclear chromosomes and was the most significantly enriched value in the entire data set (Figure 1F and G). Additionally, the average sequence length with elevated methylation in the nuclear DNA samples was 347 bp. However, nearly the entire mitochondrial genome had elevated methylation, therefore it is represented as a single data point in Figure 2F. This is the longest sequence in the entire data set with methylation identified.

**Figure 2.**
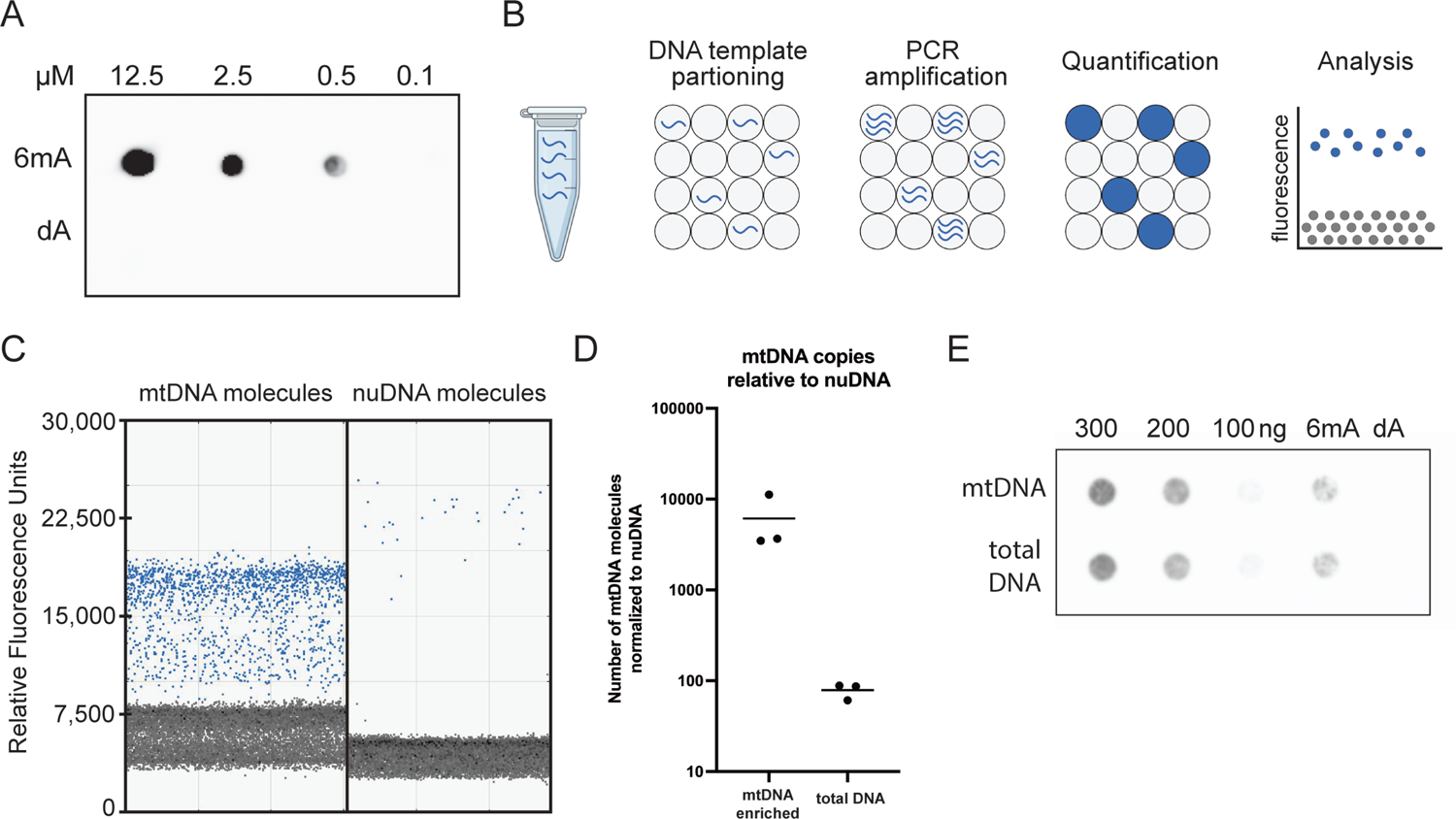
Dot blot assay shows 6mA signal in mitochondrial DNA. (A) Anti-6mA dot blot of synthetic oligonucleotides with an internal N6-methyladenosine (6mA) or without (dA) blotted at 12.5, 2.5, 0.5, and 0.1 µM. (B) Schematic of droplet digital PCR. DNA template is partitioned into individual oil droplets such that there is 1 (blue) or 0 (gray) DNA templates per droplet. The template in each droplet is amplified through PCR and a fluorescent probe is integrated. A microfluidic detector identifies fluorescent DNA positive (blue) and fluorescent DNA negative (gray) droplets which are used to quantify the number of DNA molecules. (C) Representative ddPCR assay of an enriched mtDNA sample with positive mtDNA droplets (left) and positive nuDNA droplets (right), n=3. (D) The number of mtDNA copies relative to nuDNA in the mtDNA enriched sample (left) and the total DNA enriched sample (right) from three independent biological experiments. (E) Representative anti-6mA dot blot assay of enriched mtDNA and total DNA. Three concentrations were loaded for each sample, and a positive control (6mA oligonucleotides) and negative control (dA oligonucleotides) were included.

Overall, analysis of two distinct data sets show evidence of mtDNA 6mA methylation. The fraction of methylated adenines is higher in mtDNA than nuclear DNA. There were more low-level methylation sites detected in mtDNA, but this may be due to a greater sensitivity for detecting low level sites because of higher sequence coverage. Filtering low level sites from the data showed that the average methylation level at a 6mA site is comparable in mtDNA.

### Dot blot assay shows 6mA signal in mitochondrial DNA

We next sought evidence of 6mA in mtDNA using antibody-based approaches. To validate the specificity of our anti-6mA antibody, we assessed its ability to detect decreasing amounts of an N6-methyladenosine (6mA) containing vs unmethylated (dA) oligonucleotides that are otherwise identical. The anti-6mA antibody was able to detect 6mA signal in the methylated oligo at 0.5 µM or greater (Figure 2A). No 6mA signal was detected in the un-methylated oligo sample, even at the highest concentration. This data shows that the anti-6mA antibody can discriminate 6mA from dA with high specificity.

We then used this 6mA-specific antibody to probe mtDNA from *C. elegans*. We first enriched for mtDNA or total DNA with differential centrifugation. Successful enrichment of mitochondria was confirmed by droplet digital PCR (ddPCR) using mtDNA and nuDNA specific primers. ddPCR is a highly sensitive quantitative PCR technique that quantifies DNA at single molecule resolution by running thousands of independent PCR reactions simultaneously on single DNA molecules sequestered in individual oil droplets (Figure 2B). A fluorescent probe is integrated during PCR amplification and the number of fluorescent positive droplets can then be counted by a microfluidic detector to quantify the number of DNA molecules that are present in a sample ^17^. The mitochondrial fraction showed a substantial enrichment of mtDNA and depletion of nuDNA relative to the total DNA control (Figure 2C and D and Supplemental Figure 2A). DNA from these enriched samples was standardized by dsDNA concentration and loaded onto a positively charged nylon membrane and probed with the anti-6mA antibody. Initially, we probed for 6mA using animals grown on methylation competent E. coli (Supplemental Figure 2B), however it is possible that bacterial 6mA can contribute to dot blot signal intensity. Therefore, we used animals grown on dam-/dcm-E. Coli that lack 6mA for our experiments. This showed methylation signal in the mtDNA enriched fraction (Figure 2E). This strongly suggests the presence of 6mA methylation in the *C. elegans* mitochondrial genome. Additionally, signal was detected in *C. elegans* nuclear DNA, which is consistent with previous reports ^10, 11^.

Our dot blot results suggest that *C. elegans* mtDNA is methylated. However, while a dot blot assesses 6mA signal it does not validate the signal source, therefore, any contaminating nucleotides containing 6mA could contribute to signal intensity. We therefore sought to design an assay that directly and specifically assesses mtDNA 6mA methylation.

### 6mA immunoprecipitation enriches for mtDNA

To address limitations of the dot blot assay, we performed methylated DNA immunoprecipitation (MeDIP) coupled to ddPCR (Figure 3A). mtDNA enriched samples were incubated with an anti-6mA antibody or control IgG antibody conjugated to magnetic beads. After incubation, mtDNA was released from the antibody-bead complex. We then quantified the number of mtDNA copies released via ddPCR with mtDNA specific primers. The highly specific mtDNA primers and sensitive ddPCR technique allowed us to overcome the technical limitations imposed by contaminating nucleotides. With this assay we are able to detect if the anti-6mA antibody enriched for mtDNA, regardless of contaminating nuDNA, RNA, or bacterial DNA.

**Figure 3.**
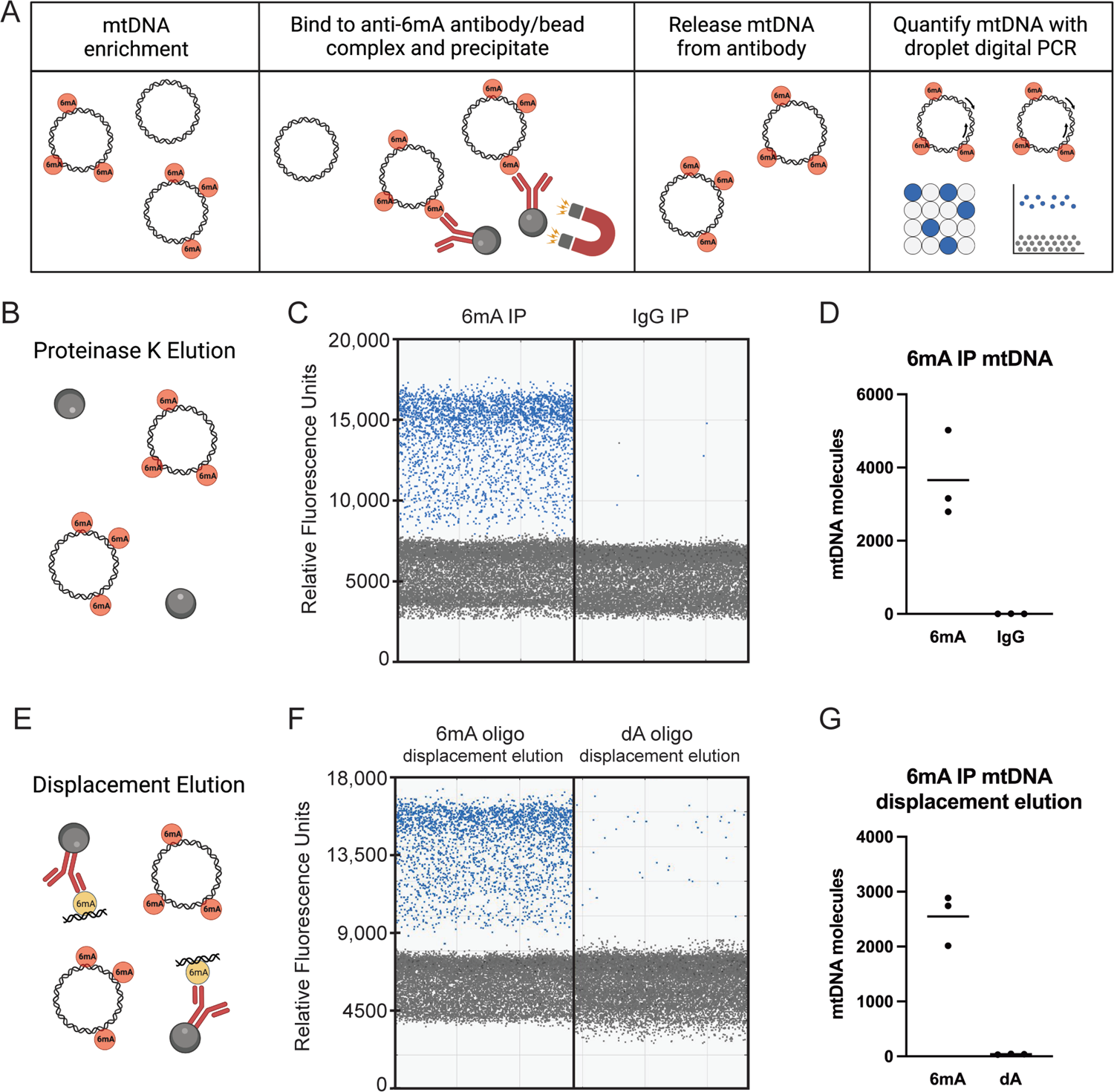
6mA immunoprecipitation enriches for mtDNA. (A) Schematic of mtDNA IP and ddPCR. mtDNA was enriched via differential centrifugation. mtDNA was then immunoprecipitated with an anti-6mA antibody/bead complex, or IgG control antibody/bead complex. mtDNA was then released from the antibody/bead complex and DNA was quantified with ddPCR. (B) In one treatment group, samples were treated with Proteinase K in elution buffer for complete release of mtDNA from antibody/bead complex. (C) Representative ddPCR results of the mtDNA positive (blue) and negative (gray) droplets in the 6mA IP (left) and the IgG control IP (right). (D). The number of mtDNA molecules immunoprecipitated in the 6mA IP (left) and the IgG control IP (right) in three independent experiments. (E) In another treatment group, 6mA IP samples were competed with 6mA methylated oligonucleotides or unmethylated oligonucleotides to displace mtDNA bound to the antibody/bead complex. (F) Representative ddPCR results of the DNA positive (blue) and negative (gray) droplets in the 6mA oligonucleotide displacement elution (left) and the dA oligonucleotide displacement elution (right). (G) The number of mtDNA molecules immunoprecipitated in the 6mA oligonucleotide displacement elution (left) and the dA oligonucleotide displacement elution (right) in three independent experiments.

We found that the anti-6mA antibody significantly enriched for mtDNA (Figure 3B-D). Three independent biological replicates showed there was 1000-fold (average 3657.33 vs 3.33) more mtDNA copies in the anti-6mA IP, compared to the control immunoprecipitation using a non-specific IgG antibody.

Although the MeDIP assay demonstrated that the anti-6mA antibody was able to enrich for mtDNA, we could not rule out the possibility that it was interacting with mtDNA irrespective of 6mA. Consequently, we designed a competitive elution whereby 6mA IP samples were eluted from the antibody bead complex either by incubation with 6mA methylated oligonucleotides (6mA) or un-methylated oligonucleotides (dA) (Figure 3E). If the anti-6mA antibody enriched for mtDNA by binding to 6mA, then competition with 6mA oligonucleotides should displace mtDNA from the antibody-bead complex. Further, if this interaction is 6mA specific, mtDNA should not be released following the addition of an unmethylated oligonucleotide. 6mA competitive elution released 70% of mtDNA when compared to complete elution with Proteinase K while dA competitive elution released only 1% (Figure 3F, 3G). Taken together, these data show that anti-6mA IP enriches for mtDNA through 6mA specific interaction, strongly suggesting that mtDNA is 6mA methylated.

### 6mA distribution within the mitochondrial genome varies between coding and non-coding, regulatory regions

Convinced that mtDNA is methylated based on initial bioinformatic analysis, dot blot, and MeDIP ddPCR 6mA enrichment, we next sought to characterize how 6mA is distributed throughout the mitochondrial genome. We first mapped 6mA sites across the genome by the percent methylation at each unique 6mA site using SMRT sequencing data (Figure 4A). This showed both high methylation and low methylation sites throughout the coding region of the genome. Koh et al., 2018 identified that 6mA was asymmetrically distributed on the heavy strand of mtDNA in cultured human cells, so we began by assessing the strand distribution of methylation sites ^13^ (Figure 4B). mtDNA methylation sites on the heavy strand have a slightly higher average methylation percentage, 33.6%, than the light strand, 31.6%. There were fewer total methylation sites on the heavy strand, but there are also fewer adenines on the heavy strand so the 6mA:dA ratio was similar – 8.28% and 8.14% on the heavy and light strands, respectively.

**Figure 4.**
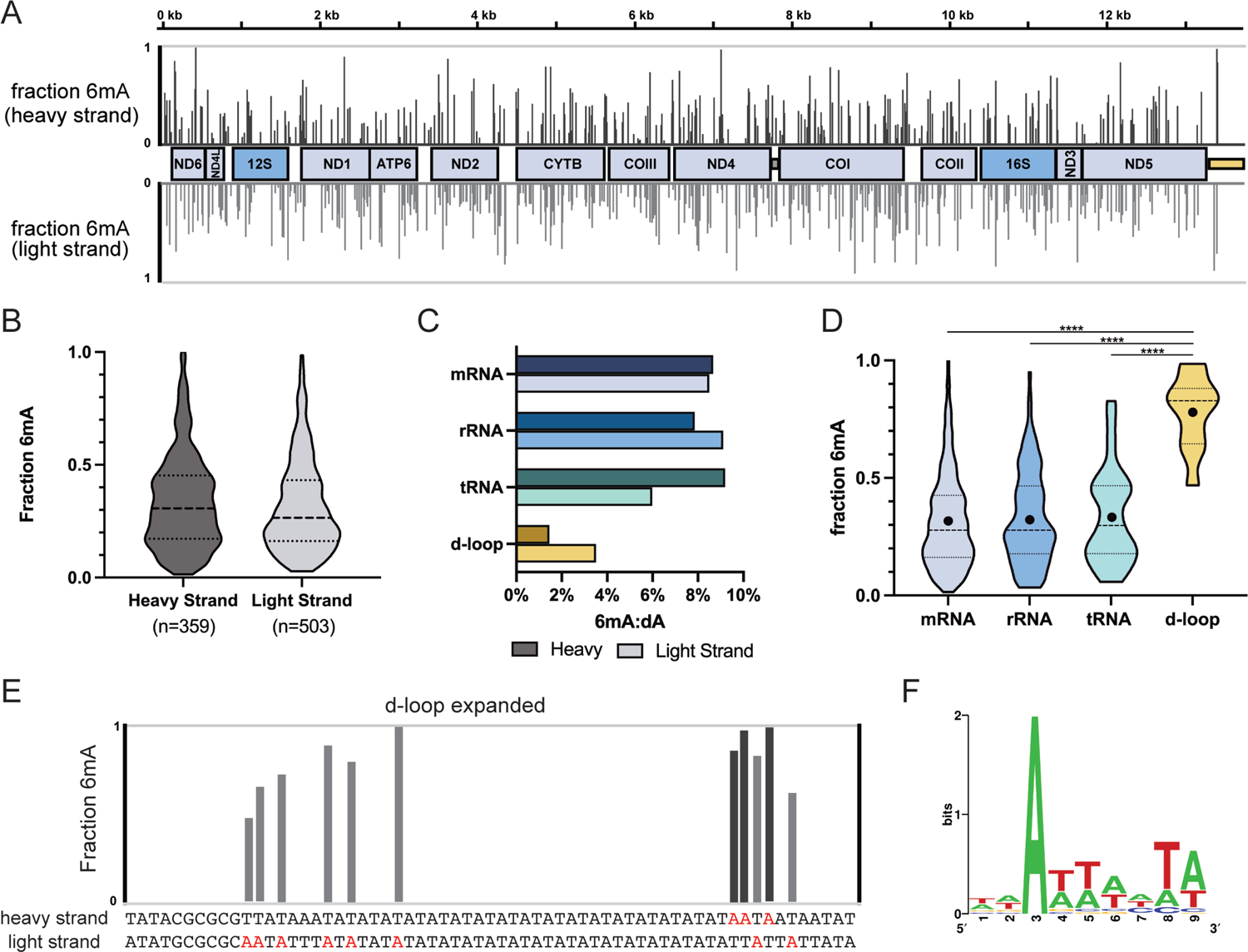
6mA distribution within the mitochondrial genome. (A) Distribution and methylation level of 6mA methylation sites in mtDNA on the heavy (dark gray, upper) and light (light gray, lower) strands. Methylation levels at a given adenine position are represented from 0 to 1, with 0 being 0% of reads were methylated at that site and 1 being 100% of reads are methylated at that site. (B) Level of 6mA methylation on the heavy (n=359, mean=33.6%) and light strand (n=503, mean=31.6%). (C) Ratio of 6mA sites to total adenines within mRNA, rRNA, and tRNA coding regions and the d-loop. Data is further stratified by heavy (darker shade) or light strand (lighter shade). (D) Methylation levels within mRNA, rRNA, tRNA coding regions and the d-loop. Mean values are denoted with a black dot and 25^th^, 50^th^, and 75^th^ distribution percentiles are marked with dashed lines. Comparisons between groups were made with an ordinary one-way ANOVA. mRNA n=662 and mean=31.5%, rRNA n=108 and mean=32.6%, tRNA n=70 and mean=33.5%, and d-loop n=11 and mean=78.2%. (E) Expanded view of methylation of the d-loop. Methylation levels are plotted at individual adenine sites in the first 100 bp of the d-loop. (F) A sequence motif enriched in at sites of methylation was detected using STREME ^25^.

We then compared the ratio of adenine methylation sites to total adenine (6mA:dA) for mRNAs, rRNAs, tRNAs, and the major non-coding region (akin to the mammalian d-loop) (Figure 4C). The fraction of adenines that are methylated is similar for mRNA, rRNA, and tRNA, with approximately equal distribution on both the heavy and light strand of mRNA and rRNA genes. Interestingly, the d-loop had the lowest ratio of 6mA:dA sites; only 11 methylation sites were identified in the very AT rich region. The 2.5% of methylated adenines found in the d-loop is substantially lower than the 8.2% average observed across the mitochondrial genome.

We then compared the percent methylation per site for mRNAs, rRNAs, tRNAs, and the d-loop. The distribution of percent methylation at 6mA sites was similar for mRNA, rRNA, and tRNA and the average was 32% (Figure 4D). Strikingly, the d-loop had a significantly (p < 0.0001) higher percent methylation, averaging 78%, at 6mA sites compared to mRNA, rRNA, and tRNA encoding regions. Additionally, all the methylation sites within the d-loop are clustered within the first 100 bp (Figure 4E). There is a cluster of 6 sites on the light strand, a space of 14 repeating ATs, and then a cluster of methylation sites on both the heavy and the light strands. These are the only sites within the d-loop that methylation was detected at, as the remainder of the d-loop is unmethylated. A weak motif is also associated with methylation (Figure 4F), but it is not clear if this motif is biologically significant. This unique distribution of 6mA between coding and non-coding regions suggests that 6mA in the mitochondrial genome is unlikely to be the result of random incorporation of methylated bases. Instead, it is likely that mtDNA is methylated through a regulated biological process.

### mtDNA 6mA levels are dynamically regulated in response to the mitochondrial stressor antimycin

In addition to characterizing mtDNA methylation in depth, we wanted to assess if mtDNA 6mA levels are dynamically regulated. To assess if *C. elegans* mtDNA methylation levels can change, we analyzed MeDIP-seq data from animals that had experienced mitochondrial stress ^11^. Animals in Ma et al., 2019 were treated with antimycin, a potent electron transport chain inhibitor that targets complex III, and 6mA MeDIP was performed followed by Illumina sequencing. This was compared to control animals that were not antimycin treated, as well as control groups that used a non-specific IgG within each treatment group. Our analysis of these data show that antimycin treatment increased the amount of mtDNA pulled down by the anti-6mA antibody at several points of the mitochondrial genome (Figure 5A). Five of these sites were statistically significant for 6mA enrichment and were within the genes encoding for *nduo-6*, *ctb-1*, *ctc-3*, *ctc-1*, and *nduo-5*. Overall, these data suggest that antimycin increased 6mA in the mitochondrial genome.

**Figure 5.**
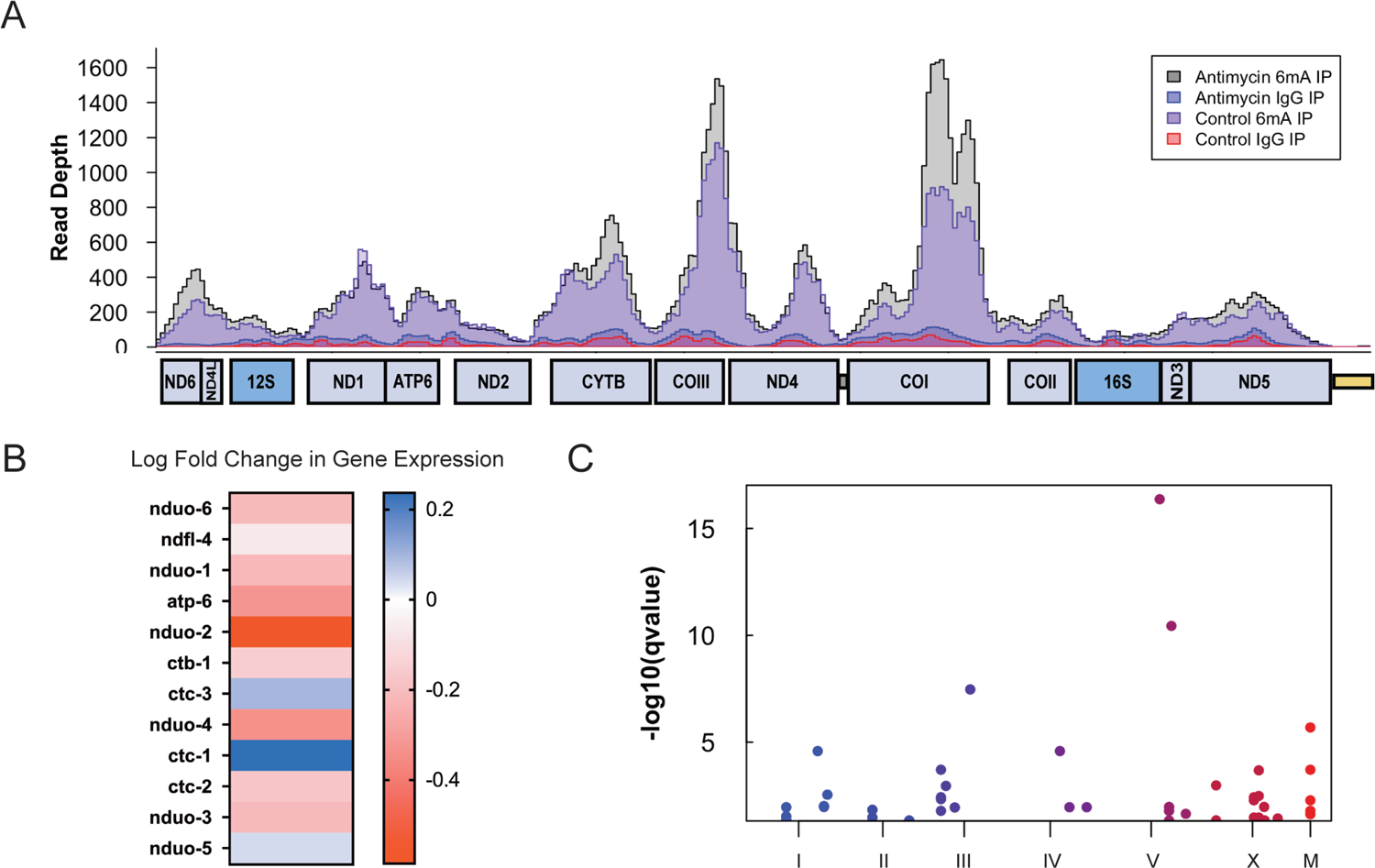
mtDNA 6mA levels are dynamically regulated in response to mitochondrial stressor antimycin. (A) MeDIP read depth throughout the mitochondrial chromosome in antimycin treated 6mA IP (black), control 6mA IP (purple), antimycin treated IgG IP (blue), and control IgG IP (red) samples. (B) Log2 fold-change in mtDNA transcript expression in antimycin treated animals compared to control animals. Color gradient is Log2 fold-change from positive fold-change (blue), to no change (white), to negative-fold change (red). (C) Significantly enriched sequences in the antimycin treated 6mA IP compared to the control 6mA IP using MACS2. The -log10 of the qvalue for each enriched sequence is plotted as a dot. ChrI n=9, ChrII n=4, ChrIII n=3, ChrIV n=4, ChrV n=7, ChrX n=4, ChrM n=10.

Although we observe a change in methylation levels, it isn’t yet clear what regulatory role increased methylation may have. One possibility is regulation of mtDNA gene expression. Notably, most mtDNA encoded transcripts had decreased expression in antimycin treated animals. However, ctc-1, ctc-3, and nduo-5 had a slightly increased expression (Figure 5B). Interestingly, these genes with increased transcripts account for three of the five sites of increased methylation.

Further, these five sites of methylation enrichment represent a disproportionate effect on mtDNA compared to nuclear DNA (Figure 5C). The nuclear chromosomes each had less than ten enriched 6mA sites, which is surprising because each chromosome is several megabases long, compared to the approximately 14kb mitochondrial genome. This indicates that antimycin has an outsized effect on 6mA in the mitochondrial genome compared to the nuclear genome. Taken together, these data suggest that mtDNA 6mA methylation can be dynamically regulated in response to mitochondrial stress.

### mtDNA 6mA levels are increased in NMAD-1 loss of function animals

We were further interested in identifying genetic perturbations that can alter methylation levels, including changes in potential methyltransferases or demethylases. Intriguingly, Wang et al. reported that NMAD-1, a protein that contains a DNA oxidative demethylase domain and is an ortholog of human ALKBH4, directly binds to the mitochondrial single stranded binding factor MTSS-1 ^18^, a protein known to be exclusively localized to mitochondria ^19^. This suggests that NMAD-1 may have a mitochondrial regulatory role. The link between NMAD-1 and mitochondrial regulation is further strengthened by evidence of an ATFS-1 binding site near *nmad-1* ^20^. ATFS-1 is a crucial transcription factor that activates the mitochondrial unfolded protein response (UPR^mt^) during mitochondrial proteolytic stress to maintain mitochondrial homeostasis ^20, 21^.

Intrigued by these links, we evaluated 6mA levels in *nmad-1* loss of function animals (strain VC2552). We observed a substantial increase in 6mA in mtDNA in *nmad-1* mutant animals compared to wildtype animals (Figure 6A). To further validate increased mitochondrial methylation, we leveraged the MeDIP-ddPCR assay to quantitatively assess 6mA in *nmad-1* -/- animals which again showed increased mitochondrial methylation (Supplemental Figure 3A). These data show that loss of NMAD-1 function lead to methylation accumulation.

**Figure 6.**
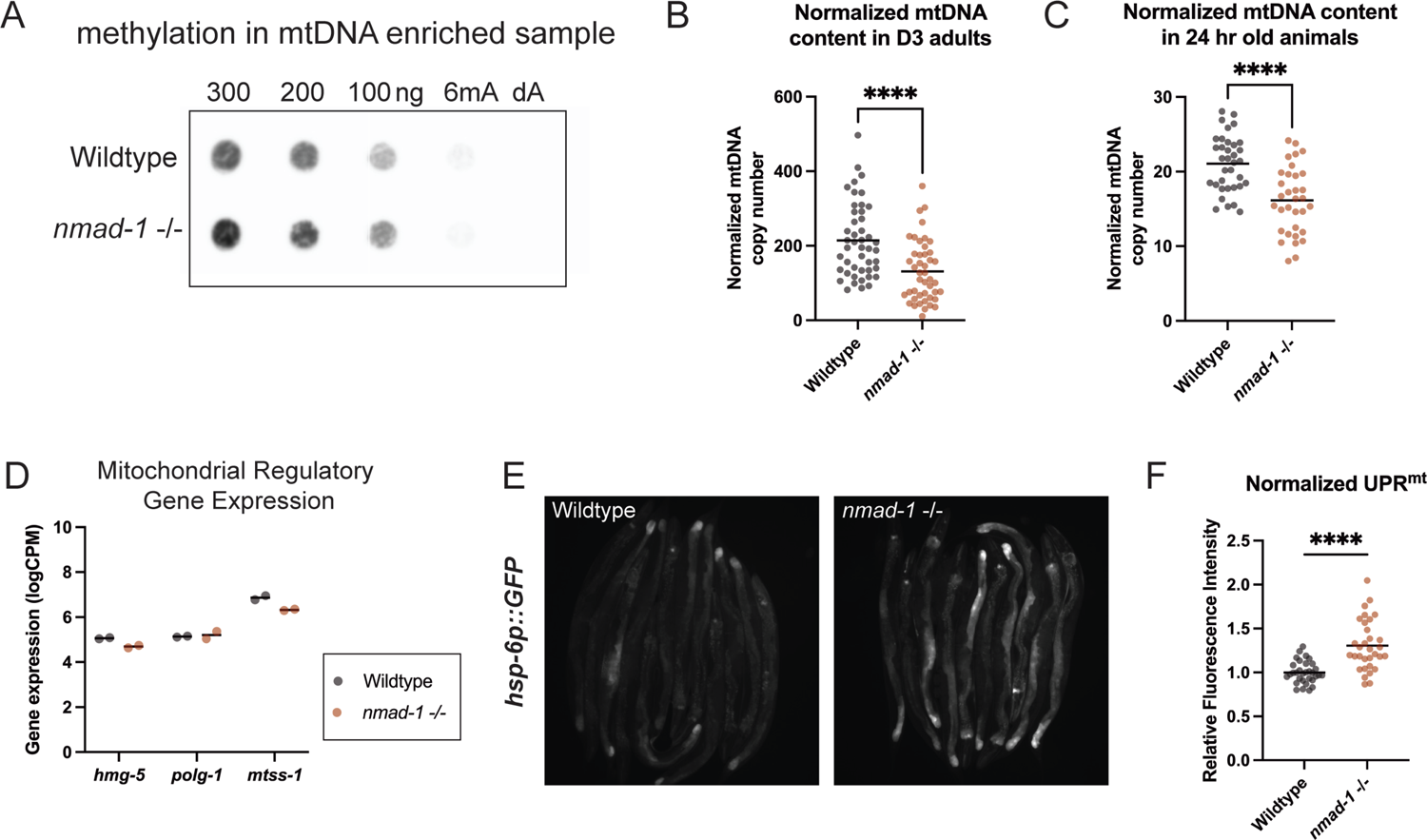
*nmad-1* loss of function animals have increased mtDNA 6mA and decreased mtDNA copy number. (A) anti-6mA dot blot of mtDNA from wildtype and *nmad-1* -/- animals with decreasing DNA concentrations, a positive control (6mA), and negative control (dA). Representative image of two independent biological replicates. (B) mtDNA copy number, normalized to nuDNA, in age synchronized day 3 (D3) adults. Wildtype n=46, *nmad-1* -/- n=45, unpaired t-test, **** indicates p-value <0.0001. (C) mtDNA copy number, normalized to nuDNA, in age synchronized animals 24 hr post embryo-lay. Wildtype n=35, *nmad-1* -/- n=33, unpaired t-test, **** indicates p-value <0.0001. (D) logCPM normalized transcript expression of *hmg-5*, *polg-1*, and *mtss-1* from wildtype (black) and *nmad-1* -/- (red) animals. Two independent replicates represented as individual data points, means shown with black bar. (E) Representative image wildtype or *nmad-1* -/- animals labeled with the *hsp-6p::GFP* reporter of UPR^mt^ (F) Relative fluorescent intensity normalized to the wildtype average of wildtype and *nmad-1* -/- animals with the *hsp-6p::GFP* reporter. Wildtype n=32, *nmad-1* -/- n=32, unpaired t-test, **** indicates p-value <0.0001.

### NMAD-1 loss of function animals have mis-regulated mtDNA copy number

Epigenetic modifications of DNA are frequently a means of regulating replication and transcription ^1, 7, 11^. We were especially interested in mtDNA replication because of the unique distribution of 6mA in the regulatory d-loop region of mtDNA, a reported region of replication initiation. To test relative replication effects, we first tested mtDNA copy number in adult animals. This revealed that *nmad-1* -/- animals have a significant decrease in mtDNA copy number compared to wildtype (Figure 6B and Supplemental Figure 3B and C). However, *nmad-1* animals are known to have germline dysregulation ^18^, and the germline contains the majority of mtDNA in an adult *C. elegans* ^22^. To rule out the possibility of germline mitochondrial mass affecting copy number, we tested copy number in larva 24 hrs post embryo-laying that have not yet begun substantial germline development (Figure 6C and Supplemental Figure 3D and E). These young animals also have a significant decrease in mtDNA copy number. Combined, our data show a correlation between increased methylation and suppressed replication.

One potential explanation for decreased mtDNA copy number would be decreased levels of mtDNA regulatory proteins. However, analysis of RNA seq data from Wang et al. demonstrates that *nmad-1* -/- animals do not show a significant change in transcript expression of the mtDNA replicative polymerase, *polg-1*, the mitochondrial packaging factor, *hmg-5*, or the mitochondrial single-stranded DNA binding protein, *mtss-1* (Figure 6D). This suggests that the mtDNA copy number effects observed are unlikely to be the result of changes in the expression of mtDNA regulatory proteins. Instead, it may reflect that 6mA alters how regulatory proteins interact with the mtDNA molecule.

We were next interested in determining if *nmad-1* -/- had consequences for mitochondrial function more broadly. Indeed, we observed that *nmad-1* animals show a modest, but significant, activation of the mitochondrial stress response UPR^mt^ (Figure 6E and F). This suggests that losing NMAD-1 function can cause proteolytic stress in mitochondria. This proteolytic stress could reflect the decrease in mtDNA copy number observed in these animals. Together, these data show that mtDNA 6mA levels increase in absence of NMAD-1 and this increase is accompanied by changes in mitochondrial function. This suggests a novel role for NMAD-1 as a mitochondrial regulator.

## Discussion

Mitochondria are semi-autonomous organelles at the heart of energy production and cell metabolism. These organelles are unique from other cellular components because they contain their own small, but essential genome that encodes for several proteins of the electron transport chain. Dysregulation of mtDNA therefore directly affects cellular energy production, so mtDNA transcription and replication must be tightly regulated. Epigenetic modifications are one way gene expression can be regulated without modifying the genome and are a powerful form of stress adaption ^11, 23^. Here we provide evidence that the mitochondria genome of the roundworm *C. elegans* is 6mA methylated. This is a novel route by which mitochondrial function may be regulated.

It has long been speculated that mitochondrial DNA may be subject to epigenetic modifications. Many efforts to assess mtDNA methylation have focused on characterizing 5mC, likely influenced by the important role of 5mC in eukaryotic genomes. Although there have been reports of 5mC in the mitochondrial genome, it seems that 5mC occurs at such low levels that even if it is present, its biological significance is unclear^24^. Now, there is a renewed effort to assess mtDNA epigenetics, but with a focus on 6mA methylation.

Recently, 6mA has been characterized in the mitochondrial genome of cultured human cells ^12, 13^. In the cell culture system, methylation is distributed throughout the mitochondrial genome and can be dynamically regulated in response to mitochondrial stress. Koh et al., 2018 identified 6mA throughout the mitochondrial genome that appears to enhance the binding of mitochondrial single stranded binding protein during replication. The model put forth by Hao et al., 2020 posits that 6mA can inhibit the binding of the major mitochondrial transcription factor, TFAM, to attenuate transcription and suppress oxidative phosphorylation during mitochondrial stress. Together, these data suggest that 6mA has an important role in mtDNA regulation, however more experiments are needed to understand the mechanisms with high granularity. Further, it is currently unclear whether mtDNA 6mA signatures can be transmitted across generations as epigenetic information or if 6mA has a role in mitochondrial diseases. These investigative efforts would be enhanced by the ability to study 6mA in a genetically tractable model organism, such as *C. elegans*. Here we have demonstrated that *C. elegans* mtDNA contains 6mA.

We show that there is 6mA in the mitochondrial genome of *C. elegans* using multiple techniques. By analyzing SMRT sequencing data, we identified that adenine in mtDNA is methylated at a higher frequency than adenine in nuclear DNA in a wildtype background. We also analyzed MeDIP-sequencing data that confirmed 6mA is present in mtDNA at high levels. The distribution difference of 6mA in mtDNA vs other chromosomes suggests it may have distinct roles in the mitochondria and nucleus and may be regulated by distinct pathways. Additionally, we detected many low-level methylation sites in mtDNA. It is currently not clear if these sites are biologically significant. It is possible that methylation levels at unique bases are low because 6mA may be transient in nature. Rather than marking a single location to indicate continuous signaling, 6mA may play a more dynamic regulatory role.

Additionally, we directly assessed mtDNA 6mA by leveraging a highly specific anti-6mA antibody. These experiments provide strong and direct evidence that mtDNA is 6mA methylated. Further, we analyzed the SMRT sequencing distribution of 6mA in mtDNA at single base resolution and find that methylation is distributed throughout the genome with approximately equal distribution in mRNA, tRNA, and rRNA encoding regions. Surprisingly, there are very few methylation sites in the d-loop, a very AT rich regulatory region. However, the few sites that are methylated are methylated at very high levels.

Importantly, we show that, not only does mtDNA contain 6mA but, 6mA can be dynamically regulated. This points to a functional role of methylation, which would be an effective means of mediating a stress response. Indeed, we find that mtDNA methylation levels can be modified in response to mitochondrial stress. In MeDIP experiments, antimycin treatment increased 6mA enrichment for mtDNA. This suggests that 6mA can respond to mitochondrial perturbation and supports the hypothesis that 6mA may regulate mtDNA transcription or translation.

Due to the role of epigenetics in nuclear DNA, it is possible that epigenetic modifications to the mitochondrial genome regulate transcription. The unique 6mA signature in the d-loop could mark the initiation or termination of transcription. A potential link between 6mA and transcription is supported by our observation that antimycin treated animals have increased methylation and an increased number of transcripts. However, it’s not clear if there is a direct, causal link between methylation and transcription nor the mechanisms by which methylation would alter transcription. Further studies will be necessary to definitively determine if methylation is serving as a transcription inhibitor or activator.

We were also able to successfully identify a genetic perturbation that can increase mtDNA methylation levels. Knockout of the *nmad-1*, a gene that contains a demethylase domain, increased methylation detected in mtDNA. This increase in mtDNA methylation was associated with a stark change in mtDNA copy number, in both adults and young animals, and a modest activation of UPR^mt^. However, it isn’t yet clear whether NMAD-1 affects mtDNA by acting as a mtDNA demethylase or if alters methylation levels indirectly. NMAD-1 has been reported to alter methylation levels in total DNA, so it is possible *nmad-1* loss of function changes nuclear transcriptomic programming which secondarily increases mtDNA methylation. Notably though, *nmad-1* loss of function animals do not have a change in transcription levels of *polg-1, mtss-1, or hmg-5*. This does not, however, exclude the possibility of changes at the protein level. Ultimately, *nmad-1* is an interesting candidate of mitochondrial regulation and warrants further investigation of potential direct or indirect means of affecting mtDNA.

Together, these experiments support that 6mA in the mitochondrial genome of *C. elegans* has a functional role. Our findings will allow for further study of mtDNA 6mA and its regulatory mechanisms in an *in vivo* model system. This research also raises several avenues of future research, especially for exploring the potential regulation of mtDNA transcription and replication by 6mA. Finally, it will be interesting to determine whether mtDNA 6mA signatures can be inherited and what information this confers. *C. elegans* is poised as the ideal model system to address this fundamental research gap.

## Author Contributions

Conceptualization, L.K.G., J.P.H., M.R.P.; Formal Analysis, L.K.G., T.J.H., J.P.H.; Investigation, L.K.G., J.P.H., S.H.S., M.R.C., E.M.; Resources, M.R.P; Writing – Original Draft, L.K.G.; Writing – Reviewing & Editing, L.K.G., J.P.H, T.J.H., M.R.P.; Visualization, L.K.G., J.P.H., T.J.H.; Supervision, M.R.P; Funding Acquisition, L.K.G. and M.R.P.

## Acknowledgements

This work was generously supported by funding provided to M.R.P. from NIH project Grants R01 GM123260 and R35 GM145378 and Discovery Award (PR170792) from the Department of Defense’s Congressionally Directed Medical Research Program, and was further funded by support to L.K.G. and J.P.H. from the Training Program in Environmental Toxicology (NIH Grant T32ES007028). L.K.G. is also generously supported by the National Defense Science & Engineering Graduate Fellowship, United States. Droplet digital PCR to quantify mtDNA and nuDNA levels was performed through the Vanderbilt University Medical Center’s Immunogenomics, Microbial Genetics and Single Cell Technologies core. Some strains were provided by the CGC, which is funded by NIH Office of Research Infrastructure Programs (P40 OD010440).

## STAR Methods

### RESOURCE AVAILABILITY

#### Lead Contact

Further information and requests for resources and reagents should be directed to the lead contact, Maulik R. Patel.

#### Materials Availability

This study did not generate any new unique reagents.

#### Data and Code Availability

Previously published data that were analyzed here are available under the GEO accession codes GSM1637798 (SMRT sequencing)^10^, GSE118268 (MeDIP sequencing and antimycin RNA sequencing)^11^, and GSE112488 (*nmad-1 -/-* RNA sequencing)^18^. Any additional information required to reanalyze the data reported in this paper is available from the lead contact upon request.

## METHOD DETAILS

### Nematode Growth

*C. elegans* strains were maintained on Nematode Growth Medium (NGM) plates seeded with OP50 *E. coli*, unless stated otherwise.

### Mitochondria Isolation

Mitochondrial isolation was performed according to Ahier et al. with modifications ^26^. In brief, worms were harvested from at least two, densely populated, 150 mm p-plates seeded with *dam^-^/dcm^-^ E. coli* (NEB) to yield a worm pellet of ∼2 mL. Nematodes were washed off the plates by rinsing repeatedly with ∼10 mL M9 and collected in a 50 mL conical tube. Worms were pelleted by centrifugation at 450 x g for 3 min. The supernatant was discarded, worms were washed three times with 15 mL of M9 buffer, and pelleted at 450 x g. Worms were then washed once with 15 mL of ddH_2_O and the supernatant was discarded. The sample was then resuspended in 3 mL of ice-cold MIB-PI (50 mM KCl, 110 mM mannitol, 70 mM sucrose, 0.1 mM EDTA (pH 8.0), 5 mM Tris-HCl (pH 7.4) supplemented with Protease Inhibitor cocktail (Roche, #04693159001) and transferred to a 7 mL Dounce tissue grinder (KIMBLE KONTES, 885302-0007). The worms were disrupted using 40 gentle strokes performed with pestle B. The disrupted worm suspension was transferred to a clean 15 mL conical tube and centrifuged at 4 C at 200 x g for 5 min to pellet worm carcasses and the supernatant was transferred to a clean 15 mL conical tube. The solution was then centrifuged at 800 x g for 10 min to pellet nuclei and whole cells. This was repeated 3 times, each time transferring the supernatant to a fresh conical tube. The supernatant was then evenly split into 1.7 mL microcentrifuge tubes and centrifuged at 12,000 x g at 4 C for 30 min to pellet mitochondria. The supernatant was discarded. For IP experiments, the mitochondrial and nuclear pellets were each resuspended in 50 µL nuclease free H_2_O and snap frozen in liquid nitrogen. Prior to dot blot, mitochondrial and nuclear pellets were each were resuspended in 150 µL TE for DNA extraction.

### DNA extraction

*C. elegans* DNA isolation was adapted from Shoura et al., 2017 with modifications ^27^. The mitochondria enriched fraction and total cell fraction were used as starting material for DNA extraction. The respective cellular fractions were resuspended in 2 mL of worm lysis buffer (0.1M Tris-Cl pH 8.5, 0.1M NaCl, 50 mM EDTA pH 8.0, 1% SDS) and supplemented with 100 µL of Proteinase K 20 mg/mL (ThermoFisher, 25530049). These were mixed by inversion and incubated for 1 hr at 62 C. After incubation, samples were supplemented with 400 µL 5 M NaCl and mixed by inversion. Samples were further supplemented with 400 µL of CTAB solution (10% CTAB, 4% NaCl) and incubated for 10 min at 37 C. To that mixture, 2 ml of chloroform (Sigma Aldrich, C2432) was added and vortexed vigorously for ten seconds. Phase separation was achieved by centrifuging at 2000 x g for 10 min at room temperature. The aqueous phase was recovered and mixed with 2 mL of Phenol:Chloroform:Isoamyl Alcohol (25:24:1) saturated with 10 mM Tris, pH 8.0, 1 mM EDTA (Sigma Aldrich, P3803). This was vortexed vigorously for 10 seconds and phase separation was achieved by centrifugation at 2000 x g for 10 min at room temperature. The aqueous phase was recovered and supplemented with 0.6 volumes of −20 C isopropanol and mixed by inversion. DNA was precipitated by chilling samples at −20 C for 30 min followed by centrifugation at 13,000 x g for 5 min at room temperature. The pellet was washed twice with ice cold 70% ethanol and then resuspended in 200 µL TE. DNA was stored at 4 C until RNase treatment.

### RNase treatment

The 200 µL DNA sample was supplemented with 20 µL of RNase A (Thermo Fisher, EN053) and mixed by flicking the tube and inverting several times. The samples were incubated for 2 hrs at 37 C. The sample was then supplemented with 20 µL of 20% SDS, 10 µL of 0.5 M EDTA pH 8.0, and 20 µL of Proteinase K. The samples were incubated for 1 hr at 62 C. Following incubation, samples were supplemented with 40 µL 10 M ammonium acetate and mixed. The DNA was extracted with an equal volume of Phenol:Chloroform:Isoamyl Alcohol saturated with TE pH 8.0. The sample was mixed vigorously by vortexing and phase separation was achieved by centrifuging 5 min at 5000 x g. The aqueous phase was recovered and was extracted again with an equal volume of chloroform. The sample was mixed vigorously by vortexing and phase separation was achieved by centrifuging 5 min at 5000 x g. DNA was precipitated by adding 2 volumes of −20 C 100% ethanol. The sample was mixed by inversion several times and allowed to chill at −20 C for 1 hr. The DNA was pelleted by centrifugation at 13,000 x g for 5 min at room temperature. The DNA pellet was washed twice with ice cold 70% ethanol and resuspended in 50 µL nuclease free water.

### Dot Blot

A positively charged nylon 0.45 µm membrane (Biodyne B, Thermo Fisher, 77016) was equilibrated in Milli-Q purified H_2_O for 10 min and then placed in the Bio-Dot Microfiltration Apparatus (Bio-Rad, 170-6545) according to manufacturer’s instructions. DNA concentration was measured using Qubit 3 Fluorometer and Qubit dsDNA Broad Range assay it (Thermo Fisher, Q32850) and standardized for desired concentration. DNA was denatured by heating samples for 10 min at 95 C then placed immediately on ice for 5 min. 50 µL of each sample was then loaded into the dot blot apparatus and light vacuum was applied until the sample flowed through the membrane. The membrane was air-dried for 15 min then the DNA was crosslinked to the membrane using the AutoCrosslink (1200 microjoules x100) program twice in a UV Stratalinker 1800 (Stratagene). The membrane was then blocked for 1 hr at room temperature in TBST with 5% milk. After blocking, the membrane was incubated with the primary antibody, anti-6mA (Abcam, ab151230), at 1:1000 in TBST with 5% milk overnight at 4 C. The membrane was then washed 3 times with TBST for 10 min before being incubated with the secondary antibody, 1:2000 Goat anti-Rabbit IgG HRP-linked (Cell Signaling, 7074S), for 1 hr at room temperature. The membrane was again washed 3 times for 10 min each and placed in fresh TBST. Clarity Western ECL Substrate (Bio-Rad) was applied and the immunoblot was imaged using a chemiluminescent imager (GE, Amersham Imager 600).

### Immunoprecipitation of 6mA

To prepare the antibody/bead complex, Dynabeads Protein G (Thermo Fisher, 100041) were resuspended and a 25 µL aliquot was transferred to a fresh tube for each sample. The tube was placed on a magnet and the supernatant was discarded. Five µg of anti-6mA (Abcam, ab151230) or control IgG (Cell Signaling, 2729S) was diluted in 200 µL PBS supplemented with 0.02% Tween. The antibody mixture was added to the Dynabeads and was incubated on a nutator at room temperature for 10 minutes. The suspension was placed on a magnet and the supernatant was removed, and the magnetic bead-antibody complex was resuspended in 200 µL PBS with Tween and washed by gentle pipetting. The beads were placed on a magnet and the supernatant was removed immediately prior to adding mitochondria sample.

The mitochondria enriched fraction previously isolated from worms was standardized after measuring the DNA concentration using Qubit. For each sample ∼250 ng of DNA was treated with 0.33 µL of RNase A for 30 min at 37 C. The sample was then placed in 50 µL lysis buffer (50 mM KCl, 10 mM Tris pH 8.3, 2.5 mM MgCl_2_, 0.45% Tween 20, 0.45% NP-40, and 0.01% gelatin in deionized H_2_O) supplemented with 100 µg/mL Proteinase K and incubated for 1 hr at 60 C followed by 10 minutes at 95 to inactivate the Proteinase K. After lysis, the sample was supplemented with 200 µL PBS to achieve a final volume of 250 µL. The sample was then added to freshly prepared antibody/bead complexes and mixed by gentle pipetting. The sample was nutated overnight at 4 C.

After overnight incubation, the antigen/antibody/bead complexes were washed twice with 500 mM NaCl and three times with 200 µL PBS and then transferred to a clean tube. After the final wash, the antigen was eluted from the antibody/bead complex. For complete elution, the antigen/antibody/bead complexes were incubated in 50 μL of Elution Buffer (50 mM Tris-HCl pH 8.0, 10 mM EDTA, 1% SDS) supplemented with 1 µL Proteinase K for 15 min at 65 C followed by 15 min at 90 C to inactivate Proteinase K. Alternatively, the DNA antigen was eluted through displacement elution. First, 500 µL of 10 µM 6mA methylated or unmethylated (dA) oligonucleotides were heated for 10 min at 95 C then immediately transferred to ice for 5 min to achieve single strandedness. The oligonucleotides were then added to 6mA IP sample and were incubated for 1 hr at room temperature with nutation. After displacement incubation, the sample was placed against a magnet to sequester magnetic beads and the eluted sample was moved to a fresh tube for further analysis.

### Single animal mtDNA and nuDNA copy number

Animals were age synchronized as Day 3 adults by selecting for animals at developmental stage L4 and lysing 72 hrs later. Animals were age synchronized at 24 hrs by allowing ten gravid adult animals to lay embryos over a two-hour period and then removing the adults. 24 hrs post embryo-lay the larva were lysed. Single animals were lysed in 10 µl lysis buffer (50 mM KCl, 10 mM Tris pH 8.3, 2.5 mM MgCl2, 0.45% Tween 20, 0.45% NP-40, and 0.01% gelatin in deionized H2O) supplemented with 100 µg/mL Proteinase K. Lysates were incubated for 1 hr at 60 C followed by 15 min incubation at 95 C to inactivate Proteinase K.

### Digital droplet PCR

To assess mtDNA copy number with mtDNA primers DNA was diluted to the appropriate concentration for accurate ddPCR sensitivity. DNA extracted from the mitochondria or total cell and nuclei fraction was diluted 100,000x. To assess nuDNA copy number with nuDNA primers, DNA extracted from the mitochondria or nuclear fraction was diluted 500x. To assess immunoprecipitation, eluted IP samples were diluted 100x in nuclease free H_2_O to achieve an appropriate dilution for digital droplet PCR (ddPCR). For single animal lysates, Day 3 adult samples were diluted 100x for mtDNA and were not diluted for nuDNA. Lysates from 24 hr old animals were diluted not diluted for mtDNA or nuDNA reactions.

Then, 2 µL of the appropriately diluted sample was combined with 0.25 µL 10 µM forward primer, 0.25 µL 10 µM reverse primer, 12.5 µL Bio-Rad QX200 EvaGreen Supermix, and 10 µL nuclease free H_2_O. Droplet generation and PCR amplification were performed according to the manufacturer’s protocol with a 58 C annealing temperature. Droplets were scored as positive or negative in the Bio-Rad QuantaSoft program.

### MeDIP-Seq analysis

Data were retrieved from SRA under GEO accession number GSE118268. Paired end reads were assessed for read quality using FastQC ^28^. Files were then trimmed using Trimmomatic v. 0.39 with default parameters for paired end reads ^29^. Trimmed files were aligned to the *C. elegans* reference genome WBcel235 using Bowtie-2 ^30^. Multi-mapped reads were removed with Sambamba ^31^. MACS2 was used to call peaks, measure read depth, generate bedgraphs, and make statistical comparisons ^32^. The MACS2 qvalue was calculated by adjusting the pvalue to limit false discovery.

### RNA Sequencing analysis

Data were retrieved from SRA under GEO accession number GSE118268 for antimycin RNA seq data and GSE112488 for *nmad-1 -/-* RNA seq data. Paried ends were assessed for read quality using FastQC and were trimmed using Trimmomatic V. 0.39 to remove Illumina adapters and low-quality reads with a 4:15 sliding window ^29^. Samples were then aligned to the *C. elegans* Ensembl genome and gtf file (WBcel235 gtf 108) with STAR and initial gene counts were recorded ^33^. Transcript counts that aligned uniquely were quantified using the featureCounts option in the package Subread ^34^. Samples were count per million normalized and samples were filtered to remove transcripts that didn’t have at least 0.25 CPM in at least two samples. A comparison matrix was generated and statistical comparisons were made using EdgeR feature in the package Statmod V 1.4.34 ^35^.

### Fluorescence microscopy

Live, whole animal imaging was performed using a Zeiss Axio Zoom V16 stereo zoom microscope. Animals were placed on a microscope slide with a 2% agar pad and immobilized with with ∼1 µl of 100 mM levamisole (ThermoFisher #AC187870100). A coverslip was applied and images for the same experiment were captured with identical exposure times.

### Fluorescence image analysis

Fluorescence image quantification was performed using FIJI. Total pixels and pixel fluorescence intensity were determined by tracing individual animals and summing the number of pixels in the bounds and their fluorescence intensity from 1-255. Mean fluorescence intensity was calculated for each worm and used to make comparisons between treatment groups.

## KEY RESOURCES TABLE

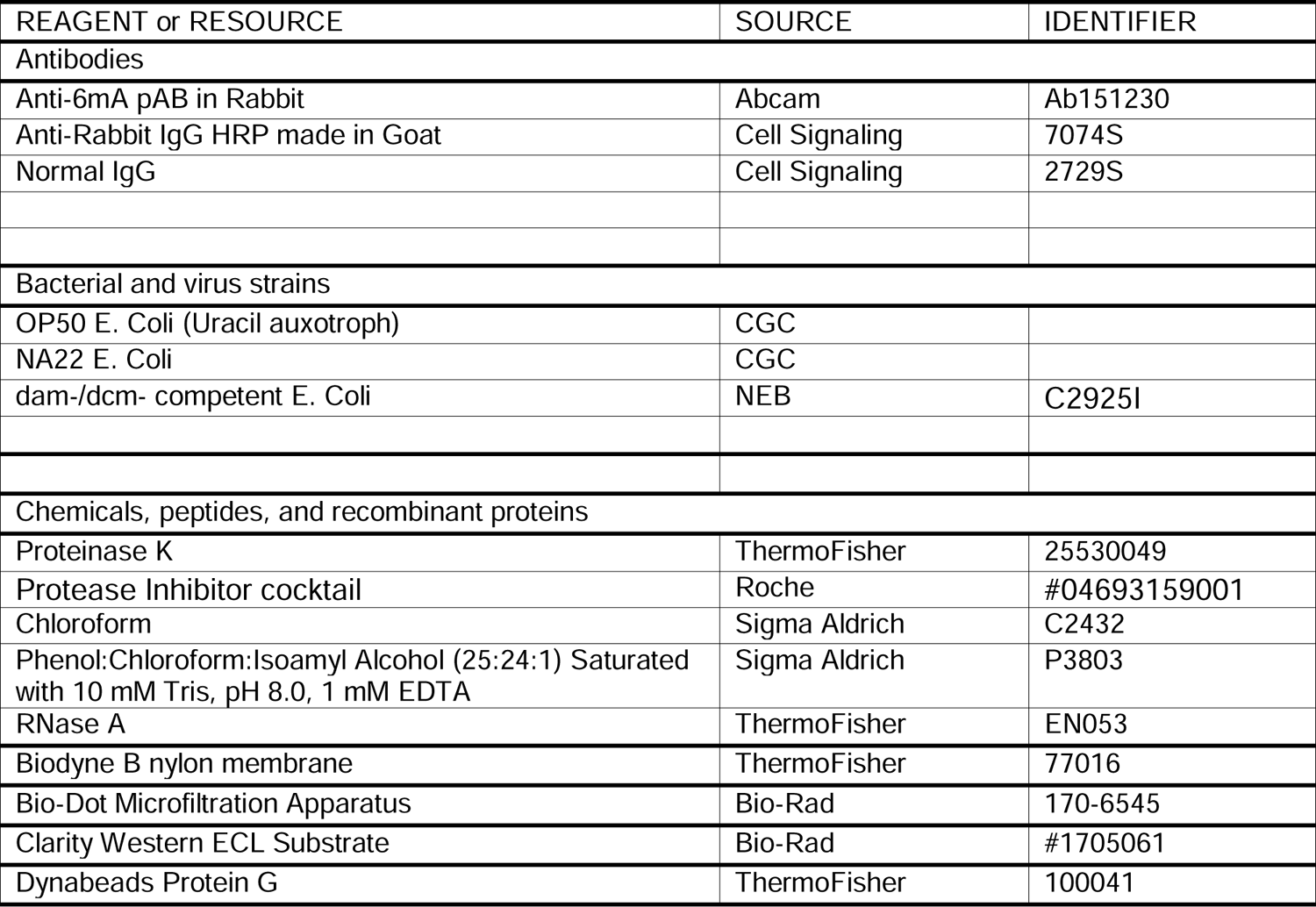
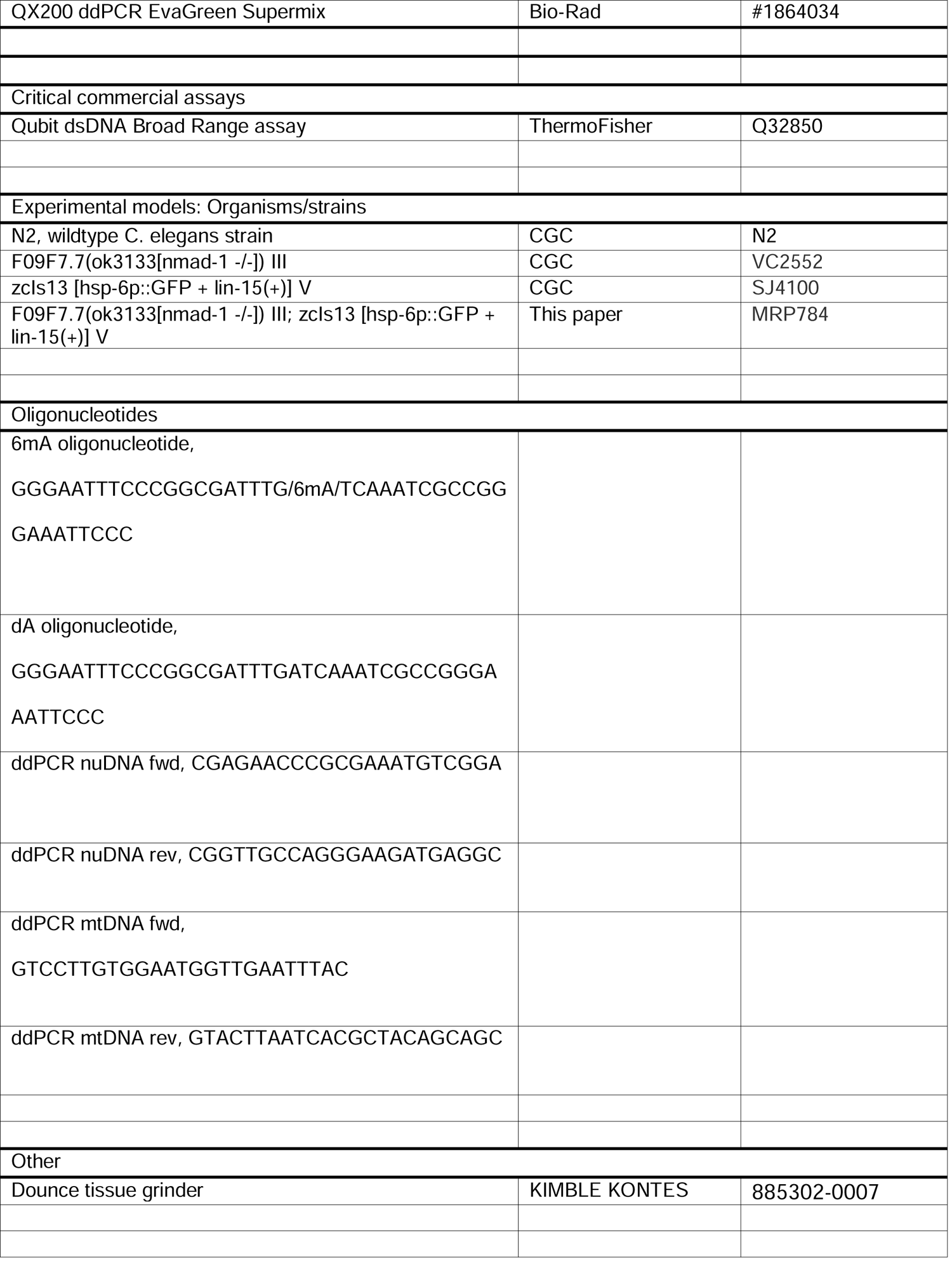

## References

1. Sánchez-Romero, M.A., and Casadesús, J. (2020). The bacterial epigenome. Nat Rev Microbiol 18, 7–20. 10.1038/s41579-019-0286-2.

2. Loenen, W.A.M., and Raleigh, E.A. (2014). The other face of restriction: modification-dependent enzymes. Nucleic Acids Res 42, 56–69. 10.1093/nar/gkt747.

3. Wion, D., and Casadesús, J. (2006). N6-methyl-adenine: an epigenetic signal for DNA– protein interactions. Nat Rev Microbiol 4, 183–192. 10.1038/nrmicro1350.

4. Fang, G., Munera, D., Friedman, D.I., Mandlik, A., Chao, M.C., Banerjee, O., Feng, Z., Losic, B., Mahajan, M.C., Jabado, O.J., et al. (2012). Genome-wide mapping of methylated adenine residues in pathogenic Escherichia coli using single-molecule real-time sequencing. Nat Biotechnol 30, 1232–1239. 10.1038/nbt.2432.

5. Fukui, K. (2010). DNA Mismatch Repair in Eukaryotes and Bacteria. Journal of Nucleic Acids 2010, e260512. 10.4061/2010/260512.

6. Stephens, C., Reisenauer, A., Wright, R., and Shapiro, L. (1996). A cell cycle-regulated bacterial DNA methyltransferase is essential for viability. Proc Natl Acad Sci U S A 93, 1210– 1214. 10.1073/pnas.93.3.1210.

7. Fu, Y., Luo, G.-Z., Chen, K., Deng, X., Yu, M., Han, D., Hao, Z., Liu, J., Lu, X., Doré, L.C., et al. (2015). N6-Methyldeoxyadenosine Marks Active Transcription Start Sites in Chlamydomonas. Cell 161, 879–892. 10.1016/j.cell.2015.04.010.

8. Mondo, S.J., Dannebaum, R.O., Kuo, R.C., Louie, K.B., Bewick, A.J., LaButti, K., Haridas, S., Kuo, A., Salamov, A., Ahrendt, S.R., et al. (2017). Widespread adenine N6-methylation of active genes in fungi. Nature Genetics 49, 964–968. 10.1038/ng.3859.

9. Zhang, G., Huang, H., Liu, D., Cheng, Y., Liu, X., Zhang, W., Yin, R., Zhang, D., Zhang, P., Liu, J., et al. (2015). N6-Methyladenine DNA Modification in Drosophila. Cell 161, 893–906. 10.1016/j.cell.2015.04.018.

10. Greer, E.L., Blanco, M.A., Gu, L., Sendinc, E., Liu, J., Aristizábal-Corrales, D., Hsu, C.-H., Aravind, L., He, C., and Shi, Y. (2015). DNA Methylation on N6-Adenine in C. elegans. Cell 161, 868–878. 10.1016/j.cell.2015.04.005.

11. Ma, C., Niu, R., Huang, T., Shao, L.-W., Peng, Y., Ding, W., Wang, Y., Jia, G., He, C., Li, C.-Y., et al. (2019). N6-methyldeoxyadenine is a transgenerational epigenetic signal for mitochondrial stress adaptation. Nature Cell Biology 21, 319–327. 10.1038/s41556-018-0238-5.

12. Hao, Z., Wu, T., Cui, X., Zhu, P., Tan, C., Dou, X., Hsu, K.-W., Lin, Y.-T., Peng, P.-H., Zhang, L.-S., et al. (2020). N6-Deoxyadenosine Methylation in Mammalian Mitochondrial DNA. Molecular Cell 78, 382–395.e8. 10.1016/j.molcel.2020.02.018.

13. Koh, C.W.Q., Goh, Y.T., Toh, J.D.W., Neo, S.P., Ng, S.B., Gunaratne, J., Gao, Y.-G., Quake, S.R., Burkholder, W.F., and Goh, W.S.S. (2018). Single-nucleotide-resolution sequencing of human N6-methyldeoxyadenosine reveals strand-asymmetric clusters associated with SSBP1 on the mitochondrial genome. Nucleic Acids Res 46, 11659–11670. 10.1093/nar/gky1104.

14. Douvlataniotis, K., Bensberg, M., Lentini, A., Gylemo, B., and Nestor, C.E. (2020). No evidence for DNA N6-methyladenine in mammals. Science Advances 6, eaay3335. 10.1126/sciadv.aay3335.

15. Flusberg, B.A., Webster, D.R., Lee, J.H., Travers, K.J., Olivares, E.C., Clark, T.A., Korlach, J., and Turner, S.W. (2010). Direct detection of DNA methylation during single-molecule, real-time sequencing. Nat Methods 7, 461–465. 10.1038/nmeth.1459.

16. Clarke, J., Wu, H.-C., Jayasinghe, L., Patel, A., Reid, S., and Bayley, H. (2009). Continuous base identification for single-molecule nanopore DNA sequencing. Nature Nanotech 4, 265–270. 10.1038/nnano.2009.12.

17. Hindson, B.J., Ness, K.D., Masquelier, D.A., Belgrader, P., Heredia, N.J., Makarewicz, A.J., Bright, I.J., Lucero, M.Y., Hiddessen, A.L., Legler, T.C., et al. (2011). High-Throughput Droplet Digital PCR System for Absolute Quantitation of DNA Copy Number. Anal. Chem. 83, 8604–8610. 10.1021/ac202028g.

18. Wang, S.Y., Mao, H., Shibuya, H., Uzawa, S., O’Brown, Z.K., Wesenberg, S., Shin, N., Saito, T.T., Gao, J., Meyer, B.J., et al. (2019). The demethylase NMAD-1 regulates DNA replication and repair in the Caenorhabditis elegans germline. PLOS Genetics 15, e1008252. 10.1371/journal.pgen.1008252.

19. Addo, M.G., Cossard, R., Pichard, D., Obiri-Danso, K., Rötig, A., and Delahodde, A. (2010). Caenorhabditis elegans, a pluricellular model organism to screen new genes involved in mitochondrial genome maintenance. Biochimica et Biophysica Acta (BBA) - Molecular Basis of Disease 1802, 765–773. 10.1016/j.bbadis.2010.05.007.

20. Nargund, A.M., Fiorese, C.J., Pellegrino, M.W., Deng, P., and Haynes, C.M. (2015). Mitochondrial and Nuclear Accumulation of the Transcription Factor ATFS-1 Promotes OXPHOS Recovery during the UPRmt. Molecular Cell 58, 123–133. 10.1016/j.molcel.2015.02.008.

21. Held, J.P., Feng, G., Saunders, B.R., Pereira, C.V., Burkewitz, K., and Patel, M.R. (2022). A tRNA processing enzyme is a key regulator of the mitochondrial unfolded protein response. Elife 11, e71634. 10.7554/eLife.71634.

22. Bratic, I., Hench, J., Henriksson, J., Antebi, A., Bürglin, T.R., and Trifunovic, A. (2009). Mitochondrial DNA level, but not active replicase, is essential for Caenorhabditis elegans development. Nucleic Acids Research 37, 1817–1828. 10.1093/nar/gkp018.

23. Mifsud, K.R., Gutièrrez-Mecinas, M., Trollope, A.F., Collins, A., Saunderson, E.A., and Reul, J.M.H.M. (2011). Epigenetic mechanisms in stress and adaptation. Brain, Behavior, and Immunity 25, 1305–1315. 10.1016/j.bbi.2011.06.005.

24. Bicci, I., Calabrese, C., Golder, Z.J., Gomez-Duran, A., and Chinnery, P.F. (2021). Single-molecule mitochondrial DNA sequencing shows no evidence of CpG methylation in human cells and tissues. Nucleic Acids Research 49, 12757–12768. 10.1093/nar/gkab1179.

25. Bailey, T.L. (2021). STREME: accurate and versatile sequence motif discovery. Bioinformatics 37, 2834–2840. 10.1093/bioinformatics/btab203.

26. Ahier, A., Dai, C.-Y., Tweedie, A., Bezawork-Geleta, A., Kirmes, I., and Zuryn, S. (2018). Affinity purification of cell-specific mitochondria from whole animals resolves patterns of genetic mosaicism. Nature Cell Biology 20, 352–360. 10.1038/s41556-017-0023-x.

27. Shoura, M.J., Gabdank, I., Hansen, L., Merker, J., Gotlib, J., Levene, S.D., and Fire, A.Z. (2017). Intricate and Cell Type-Specific Populations of Endogenous Circular DNA (eccDNA) in Caenorhabditis elegans and Homo sapiens. G3 Genes|Genomes|Genetics 7, 3295–3303. 10.1534/g3.117.300141.

28. Andrews, S. (2010). FastQC: A Quality Control Tool for High Throughput Sequence Data.

29. Bolger, A.M., Lohse, M., and Usadel, B. (2014). Trimmomatic: a flexible trimmer for Illumina sequence data. Bioinformatics 30, 2114–2120. 10.1093/bioinformatics/btu170.

30. Langmead, B., Trapnell, C., Pop, M., and Salzberg, S.L. (2009). Ultrafast and memory-efficient alignment of short DNA sequences to the human genome. Genome Biology 10, R25. 10.1186/gb-2009-10-3-r25.

31. Tarasov, A., Vilella, A.J., Cuppen, E., Nijman, I.J., and Prins, P. (2015). Sambamba: fast processing of NGS alignment formats. Bioinformatics 31, 2032–2034. 10.1093/bioinformatics/btv098.

32. Zhang, Y., Liu, T., Meyer, C.A., Eeckhoute, J., Johnson, D.S., Bernstein, B.E., Nusbaum, C., Myers, R.M., Brown, M., Li, W., et al. (2008). Model-based Analysis of ChIP-Seq (MACS). Genome Biology 9, R137. 10.1186/gb-2008-9-9-r137.

33. Dobin, A., Davis, C.A., Schlesinger, F., Drenkow, J., Zaleski, C., Jha, S., Batut, P., Chaisson, M., and Gingeras, T.R. (2013). STAR: ultrafast universal RNA-seq aligner. Bioinformatics 29, 15–21. 10.1093/bioinformatics/bts635.

34. Liao, Y., Smyth, G.K., and Shi, W. (2013). The Subread aligner: fast, accurate and scalable read mapping by seed-and-vote. Nucleic Acids Res 41, e108. 10.1093/nar/gkt214.

35. Giner, G., and Smyth, G. K., (2016). statmod: Probability Calculations for the Inverse Gaussian Distribution. The R Journal 8, 339. 10.32614/RJ-2016-024.

