## Supplementary Figures for "A role for N6-methyldeoxyadenosine in *C. elegans* mitochondrial genome regulation"

Supplemental Figure 1

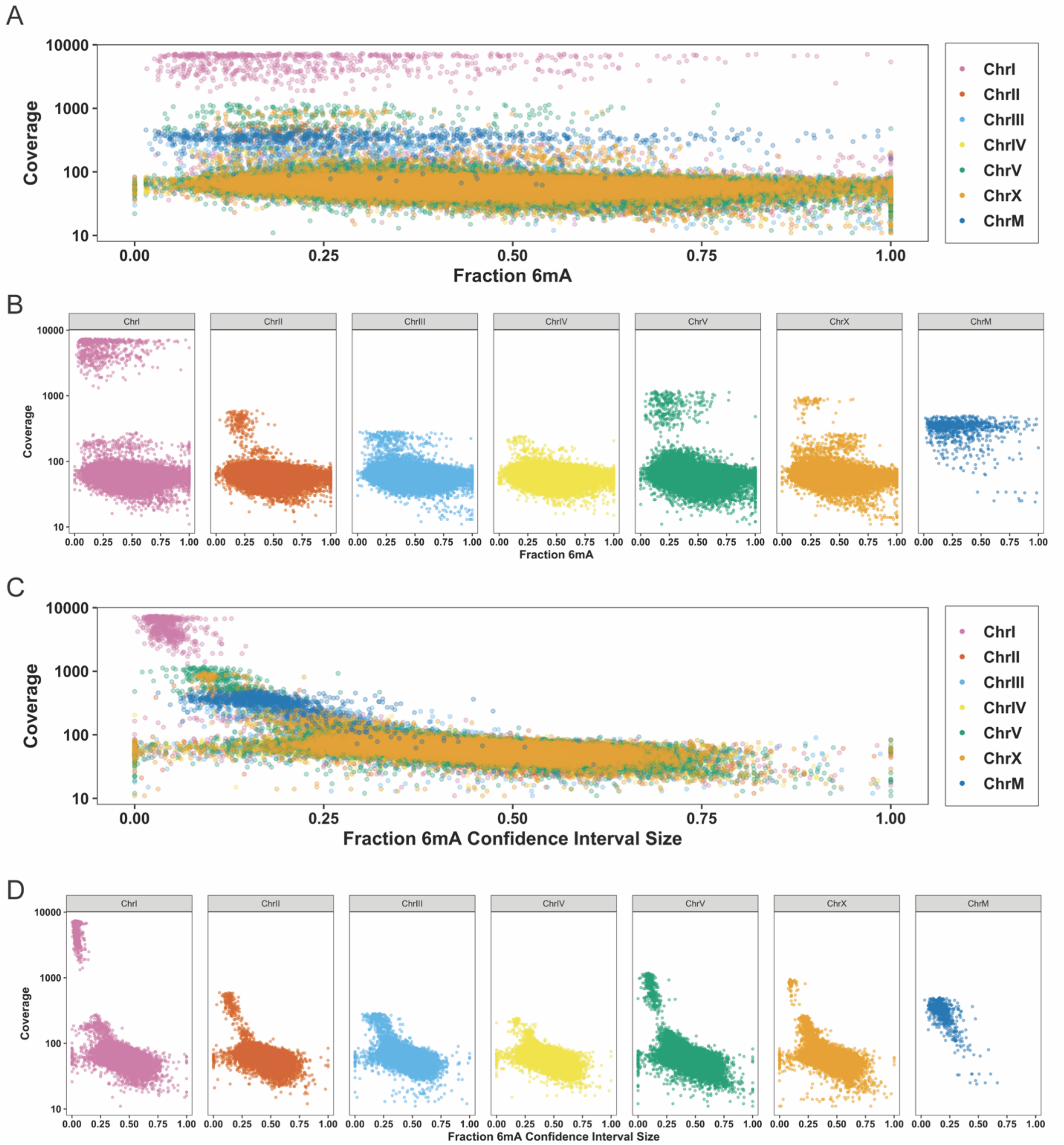

**Supplemental Figure 1.**

(A) SMRT sequencing read depth coverage compared to the fraction of methylated adenines to total adenines, stratified by chromosome. (B) SMRT sequencing read depth coverage compared to the size of the confidence interval estimating the fraction of methylated adenines to total adenines. Spearman's rank correlation was performed; p-value $< 2.2e-16$ and rho = -0.5668.

Supplemental Figure 2

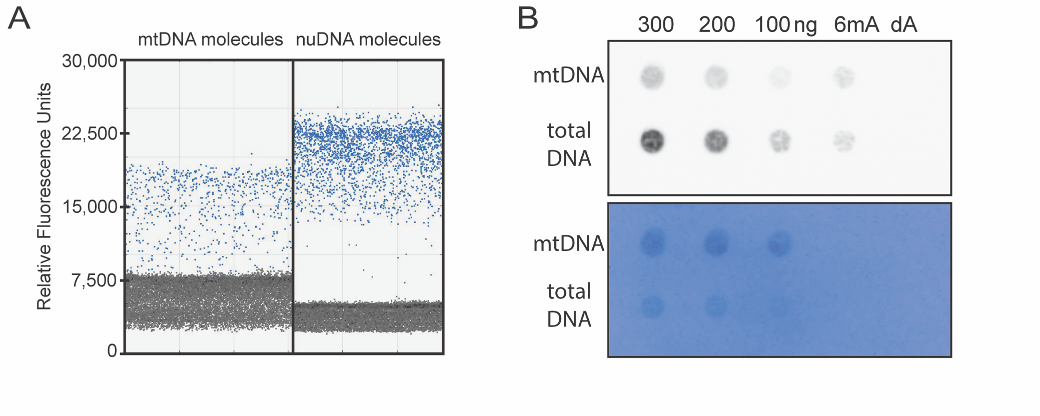


**Supplemental Figure 2.**

(A) Representative ddPCR assay of a total DNA sample with positive mtDNA droplets (left) and positive nuDNA droplets (right). (B) anti-6mA dot blot using wildtype samples showing mtDNA and total DNA 6mA content at several DNA concentrations with positive (6mA) and negative (dA) controls. Representative image of three independent biological replicates. Wildtype animals in this assay were grown in NA22 competent *E. coli*. Bottom shows methylene blue staining of membrane for total DNA.

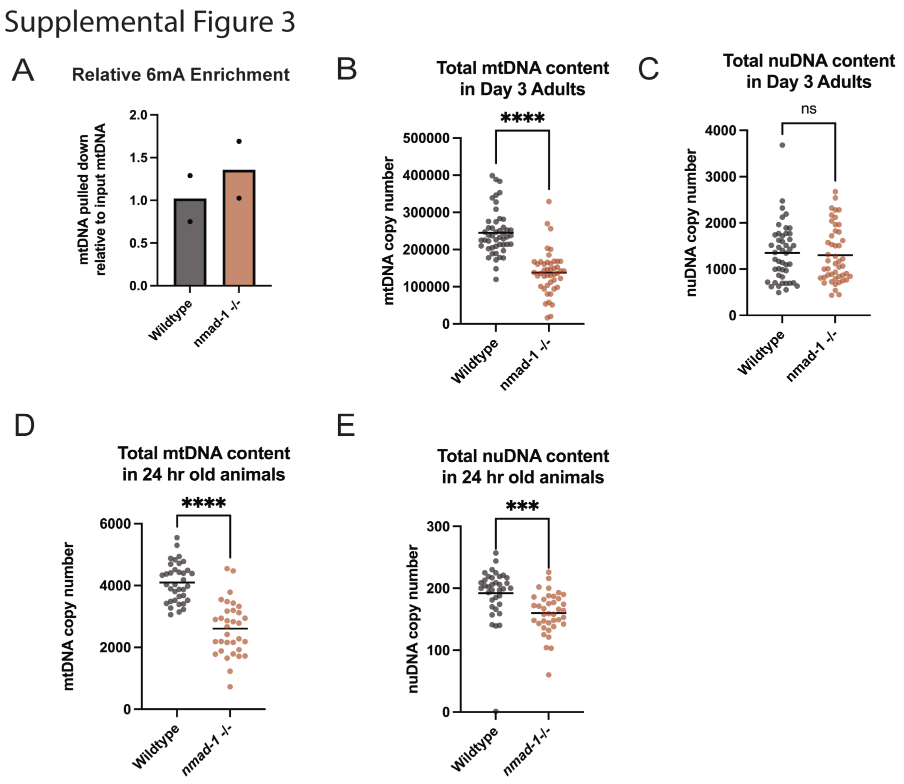


**Supplemental Figure 3.**

(A) Immunoprecipitation using anti-6mA antibody of mtDNA from wildtype and *nmad-1* -/- animals. mtDNA copy number was measured with ddPCR. The number of mtDNA copies pulled down was normalized to the input of each sample. Two independent biological replicates are shown. (B) Total mtDNA copy number in Day 3 adults in single wildtype and *nmad-1* -/- animals. Wildtype n=48, *nmad-1* -/- n=45, unpaired t-test, **** indicates p-value <0.0001. (C) Total nuDNA copy number in Day 3 adults in single wildtype and *nmad-1* -/- animals. Wildtype n=46, *nmad-1* -/- n=47, unpaired t-test, ns indicates the p-value (p-value=0.6897) is not significant. (D) Total mtDNA copy number in young animals 24 hrs post embryo-lay in single wildtype and *nmad-1* -/- animals. Wildtype n=36, *nmad-1* -/- n=33, unpaired t-test, **** indicates p-value <0.0001. (E) Total nuDNA copy number in young animals 24 hrs post embryo-lay in single wildtype and *nmad-1* -/- animals. Wildtype n=36, *nmad-1* -/- n=38, unpaired t-test, *** indicates p-value < 0.001 (p-value = 0.0006).
